# The Probability of Fusions Joining Sex Chromosomes and Autosomes

**DOI:** 10.1101/2020.03.28.010751

**Authors:** Nathan W. Anderson, Heath Blackmon

## Abstract

Chromosome fusion and fission are primary mechanisms of karyotype evolution. In particular, the fusion of a sex chromosome and an autosome has been proposed as a mechanism to resolve intralocus sexual antagonism. If sexual antagonism is common throughout the genome, we should expect to see an excess of fusions that join sex chromosomes and autosomes. Here, we present a null model that provides the probability of a sex chromosome autosome fusion, assuming all chromosomes have an equal probability of being involved in a fusion. This closed-form expression is applicable to both male and female heterogametic sex chromosome systems and can accommodate unequal proportions of fusions originating in males and females.

## Introduction

The fusion and fission of chromosomes are two of the primary mechanisms that restructure the genome into discrete chromosomes (Blackmon et al. 2019). Early on, it was recognized that both fusions and fissions might be selectively favoured because they modify linkage among loci (White 1977; Stebbins et al. 1971). In particular, the fusion of a sex chromosome and an autosome (SA-fusion) has been proposed to resolve sexual antagonism. Therefore, these fusions are predicted to be more common than autosome autosome fusions (AA-fusions) (Charlesworth and Charlesworth 1980). Limited empirical examples have shown instances where autosomes, which are enriched for sexual antagonistic loci, have recently fused with sex chromosomes (Zhou and Bachtrog 2012). For instance, a recent fusion between the X chromosome and an autosome in *Drosophila americana* is proported to have been driven by selection to reduce recombination between the sex determining locus and sexually antagonistic locus located on the autosome (McAllister 2003). Additionally, an apparent surplus in X chromosome autosome fusions in jumping spiders, *Habronattus*, is hypothesized to result from a mechanism of isolating male-beneficial sexually antagonistic alleles on the neo-Y chromosome (Maddison and Leduc-Robert 2013). Further empirical studies suggest that sexual antagonism may be common throughout the genome (Innocenti and Morrow 2010; Cheng and Kirkpatrick 2016). However, there remains significant debate on the ubiquity of sexually antagonistic variation (Kasimatis et al. 2019; Ponnikas et al. 2018). A strong measure of the frequency of significant sexually antagonistic variation across the genome would be an excess of SA-fusions relative to AA-fusions across large clades. We derive equations describing the probability of each type of fusion necessary to perform such a test.

## The Model

The probability of SA-fusions is a function of the sex chromosome system and the number of autosomes in the genome. To facilitate tests of the balance between SA-fusions and AA-fusions, we have derived a closed form expression of the probability of a SA-fusion under a null model where any chromosome is equally likely to fuse with any other non-homologous chromosome. Our result is applicable to XO, XY and multi-XY (e.g. XXY or XYY) sex determination systems and, with slight modification, to ZW systems. We ignore fusions among homologous chromosomes, including fusions that join an X and Y chromosome, because this would lead to unbalanced gametes during meiosis and, presumably, these would be non-viable. For simplicity, we first examine the case where fusions have equal probability of occurring in males and females, though we show how unequal probabilities can be accommodated. We begin with the most intuitive case, an XY sex chromosome system, and then proceed to generalize this result to more complex sex chromosome systems.

### XY System

When any two chromosomes fuse, there are 3 possibilities. The two chromosomes could both be autosomes (AA-fusion), they could both be sex chromosomes (SS-fusion), or one could be a sex chromosome and the other an autosome (SA-fusion). We will denote our three possibilities as events *AA, SS*, and *SA*, respectively. Given that a fusion has occurred, we are interested in the probability it is a SA-fusion. Or, equivalently, we are interested in the expected proportion of all fusions which are SA-fusions. Unfortunately, this proves difficult to calculate directly. We can avoid this using the complement rule. We define the probability that any given fusion is a SA-fusion as:

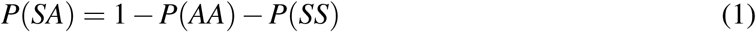

We now calculate *P*(*AA*) and *P*(*SS*) using counting. We begin with the probability of an AA-fusion, *P*(*AA*). Because we assume every chromosome is equally likely to be ‘chosen’ to fuse, we can calculate the probability that an autosome is ‘chosen’ first, *P*(*A*_1_), as the ratio of the number of autosomes to the total number of chromosomes. 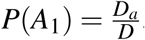, where *D*_*a*_ is the diploid autosome number and *D* is the total diploid number. The probability that the second chromosome involved in the fusion is also an autosome, *P*(*A*_2_), can be found in a similar manner. However, the first chromosome cannot be ‘chosen’ again to fuse with itself, nor can its homolog be ‘chosen’. So, the number of autosomes available to be ‘chosen’ is *D*_*a*_ − 2, the number of autosomes minus the one already chosen and its homolog. Similarly, the total number of chromosomes available is *D* − 2. Thus, 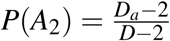, and, by independence, we have 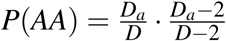. Next, we calculate the probability of a SS-fusion. Our assumption is a chromosome cannot fuse with itself, nor with its homolog. In an XY system, there are only two sex chromosomes. There is an X chromosome and either a homologous X (in females) or a homologous Y (in males). Because the sex chromosomes in an XY system are a single pair of homologs, a SS-fusion cannot occur and can be ignored. We will revisit this later in multi-XY sex chromosome systems.

Therefore, we find the probability of a SA-fusion in an XY sex chromosome system, *P*(*SA*_*XY*_), is:

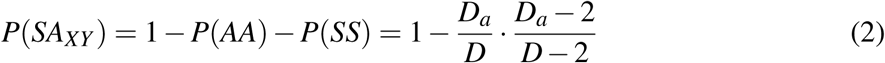

### XO System

Equation 2 does not extend to an XO system because of differences in the sex chromosome complement of males and females. In this system, males have a single X chromosome with no homolog, and females have a pair of homologous X chromosomes. The lack of a homolog in males causes males and females to have different diploid numbers and requires us to consider males and females separately.

We begin with females; following the same logic as above, we calculate the probability that an autosome is ‘chosen’ as the ratio of the number of autosomes to the total number of chromosomes present in females. 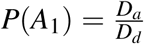, where *D*_*d*_ is the diploid number in dams. We use a subscript *s* and *d* for sire and dam when referring to sex specific values to avoid any confusion stemming from using subscript *m* and *f*. The probability that the second chromosome involved in the fusion is also an autosome can be found as the ratio of the number of autosomes available to be ‘chosen’, *D*_*a*_ − 2, and the total number of chromosomes available, *D*_*d*_ − 2. 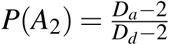. After employing independence and equation 1, we find a very familiar equation for the probability of a SA-fusion in females, *P*(*SA*_*d*_).

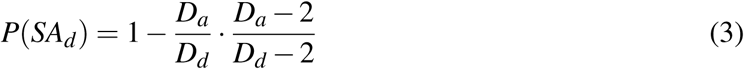

The male case, *P*(*SA*_*s*_), follows similarly and we find a nearly identical expression. The only modification required is to replace *D*_*d*_ with *D*_*s*_, the diploid number of sires, in the denominator.

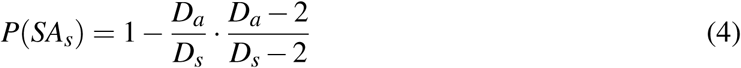

As in the XY system, we can ignore the possibility of a SS-fusion in both sexes because, in an XO sex determination system, all of an individual’s sex chromosomes are homologous.

Because we assume that males and females make equal contributions to possible fusions, we calculate the probability of a SA-fusion as the average of the probabilities that such a fusion occurs in either sex.

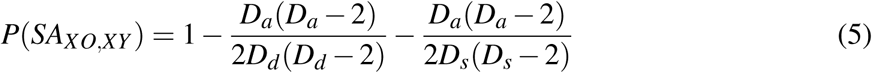

Note that in an XY system (where *D*_*s*_ = *D*_*d*_), the two fractions will combine and equation 5 will simplify into equation 2. Hence, this result is accurate for both XO and XY sex chromosome systems.

### XXY System

Recall equation 1: *P*(*SA*) = 1 − *P*(*AA*) − *P*(*SS*). In the preceding cases, we have been able to ignore the last term, *P*(*SS*). This is not the case in multi-XY systems. For example, in an XXY system females have four X chromosomes (two homologous pairs) and males have two non-homologous X chromosomes and a Y chromosome. So, in order to modify equation 5 for an XXY system, we need only find an expression for the probability of a SS-fusion in both males and females. In an XXY system, females and males have different diploid numbers, so we, again, consider the male and female cases separately.

Females in an XXY system will have four X chromosomes, two pairs of homologs, and *D*_*a*_ autosomes. We calculate the probability of a SS-fusion as the product of the probability of a sex chromosome being ‘chosen’ to fuse first, *P*(*S*_1_), and the probability of a sex chromosome being ‘chosen’ to fuse second, *P*(*S*_2_). Proceeding by counting, we calculate the probability that a sex chromosome is ‘chosen’ first, 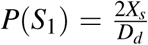, where *X*_*s*_ is the number of X chromosomes present in sires. The use of 2*X*_*s*_ in females takes advantage of the fact females always have twice as many X chromosomes as males and avoids the use of another variable for the number of X chromosomes in females. The probability that the second chromosome involved in the fusion is also a sex chromosome can be found in the same manner. The number of sex chromosomes available to be ‘chosen’ is 2*X*_*s*_ − 2, and the total number of chromosomes available is *D*_*d*_ − 2. It follows 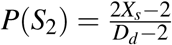. We find the probability of a SS-fusion in females is 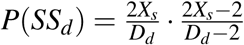. Appending this result to equation 3, we find the probability of a SA-fusion in females:

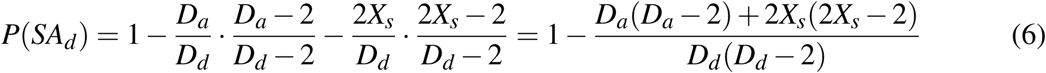

XXY males have two non-homologous X chromosomes, a single Y chromosome, and *D*_*a*_ autosomes. The Y chromosome cannot fuse with either of the X chromosomes, because of our assumption with regard to fusions of homologous chromosomes. The only possible SS-fusion is between the two non-homologous X chromosomes. We calculate the probability of a SS-fusion as the product of the probability to ‘choose’ the first X chromosome, *P*(*X*_1_), and the probability of ‘choosing’ the second X chromosome, *P*(*X*_2_). We calculate *P*(*X*_1_) as the ratio of X chromosomes to the total number of chromosomes, 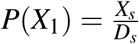. We calculate *P*(*X*_2_) as the ratio of the number of remaining X chromosomes (*X*_*s*_ − 1 only the single X chosen must be accounted for since it has no homologous X that could be chosen) and the total number of chromosomes available to fuse (*D*_*s*_ −2, every chromosome except for the X that was ‘chosen’ and the Y). Therefore, 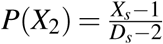 and the probability of a SS-fusion in XXY males 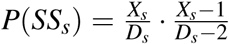. Appending this result to equation 4:

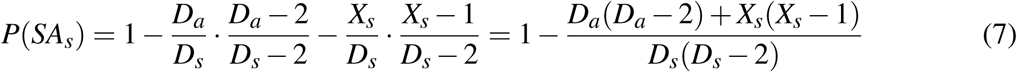

To formulate our general expression for XXY, XY and XO systems, we average the contribution from males and females and simplify.

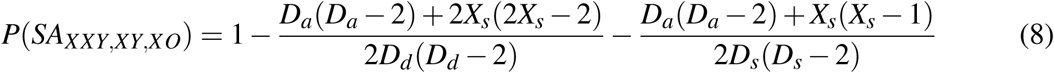

### XYY System

In an XYY system, males have a single X chromosome and two non-homologous Y chromosomes, while females have a single pair of homologous X chromosomes. The only sex chromosomes in females are an X and its homolog and there is no possibility of a SS-fusion. Recall in equation 6, the probability of both chromosomes in a fusion being sex chromosomes in a female is captured by the expression 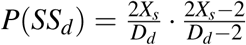. In an XYY system, *X*_*s*_ = 2 and *P*(*SS*_*d*_) = 0. Therefore, equation 6 is appropriate for females in an XYY systems as well. However, in males a SS-fusion between the two Y chromosomes is possible. As previously mentioned, we ignore the possibility of either of the Y chromosomes fusing with the X. So, the probability of a SS-fusion in males is equivalent to the probability of ‘choosing’ one Y and then the other. Proceeding by counting, we find 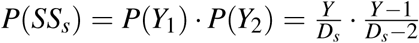 where *Y* is the number of Y chromosomes in males. Appending this to equation 4 we get:

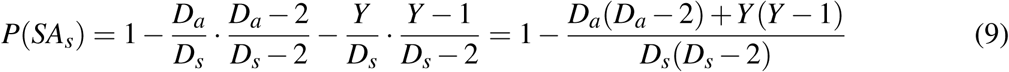

The only difference between equations 7 and 9 is *X*_*s*_ changes to *Y* in the numerator. To generate an expression that is applicable to both XXY and XYY systems we take the maximum value among *X*_*s*_ and *Y* :

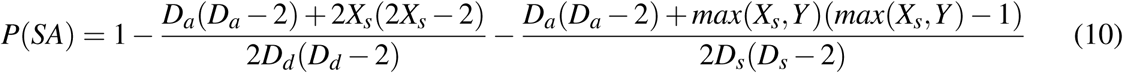

This formulation is applicable to XO, XY and multi-XY sex chromosome systems. It is quite possible that the sexes may make unequal contributions to the fusions entering a species (Pennell et al. 2015). In this case, equation 10 can be modified by the addition of a term *µ*_*d*_, representing the proportion of fusions that occur in females:

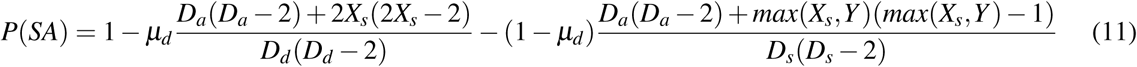

As a corollary, we are also able to derive general expressions for *P*(*SS*) and *P*(*AA*) by averaging our previous results for *P*(*SS*_*s*_) and *P*(*SS*_*d*_), and *P*(*AA*_*s*_) and *P*(*AA*_*d*_).

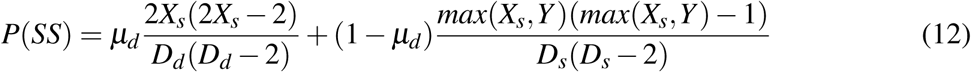

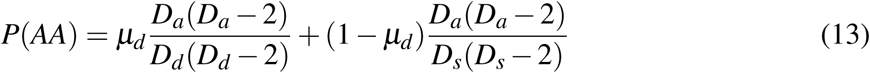

Equations 11-13 have six parameters: *µ*_*d*_, *X*_*s*_, *D*_*a*_, *Y, D*_*d*_ and *D*_*s*_. Recall, that we had eliminated one parameter, *X*_*d*_, by noting *X*_*d*_ = 2*X*_*s*_. We can eliminate two more variables by substituting *D*_*d*_ = 2*X*_*s*_ + *D*_*a*_ and *D*_*s*_ = *X*_*s*_ + *Y* + *D*_*a*_. Although illustrated for male heterogametic systems, these formulations can be converted for use in ZW sex chromosome systems as well. Taking equations 11 - 13 and exchanging *D*_*d*_ and *D*_*s*_, replacing *X*_*s*_ with *Z*_*d*_,replacing *Y* with *W*, and replacing *µ*_*d*_ with *µ*_*s*_, generates equations that will provide probabilities for ZO, ZW, and multi-ZW systems. We have provided equation 11, 12 and 13, and their ZW equivalents, as R functions in *supplemental file 1*.

## Results and Discussion

There are several cases where the derived equation, *P*(*SA*), will fail. First, in systems with UV sex chromosomes. In these systems, it is the gametophyte stage that occurs as separate males (carrying a V chromosome) and females (carrying a U chromosome) (Bachtrog et al. 2014). Second, in systems with multiple X and multiple Y chromosomes (e.g. the platypus carries 5 X and 5 Y chromosomes) our formulation will fail to provide accurate probabilities (Hsu and Benirschke 2013). However, these systems are exceedingly rare across the tree of life. Among 14,147 surveyed invertebrates just 0.4% possess these systems, and the vast majority of these (52 species) are all termites in the order Blattodea (Blackmon et al. 2017). These sex chromosome systems are equally rare in mammals where they are restricted to two species in Monotremata (Ashman et al. 2014).

The need for a quantitative null model of the probability of SA-fusions is illustrated by examining the expected probability of SA-fusions across a range of observed chromosome numbers and sex chromosome systems. In figure 1, we show when the autosome number is small, a large proportion of fusions are expected to be SA-fusions even under a null model which assumes they are not selectively favored. In fact, for the XY sex chromosome system the probability of a given fusion being an SA-fusion does not drop below 25% until the diploid autosome count is greater than 16. In systems with XXY sex chromosomes, the case is even more extreme. The probability of SA-fusion does not drop below 25% until the diploid autosome count is greater than 22. Therefore, evaluating the proportion of SA-fusions and determining whether there is evidence for positive selection on these fusions can only be accomplished in light of a quantitative null model which accounts for chromosome number and sex chromosome system. In a recent study of jumping spiders, *Habronattus*, the large disparity between the number of SA-fusions (8-15) and AA-fusion (1) and SS-fusions (1) all in a system with 26 autosomes is presented as evidence that SA-fusions are being favored (Maddison and Leduc-Robert 2013). The intuition that this pattern is unlikely can be rigorously tested with our null model. Using our equations 11-13, and a multinomial distribution, we are able to calculate the exact empirical p-value of having observed eight or more SA-fusions out of a total of 10 fusions. We assume an XXO sex chromosome system and a diploid autosome count of 26 (this karyotype was the most common in the ancestral state estimation performed in the study). 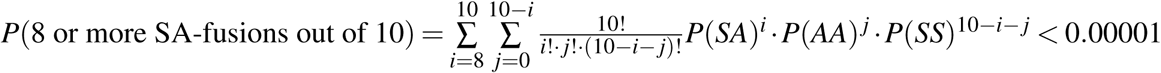. This confirms that *Habronattus* spiders do in fact have an excess of SA-fusions.

**Figure 1:**
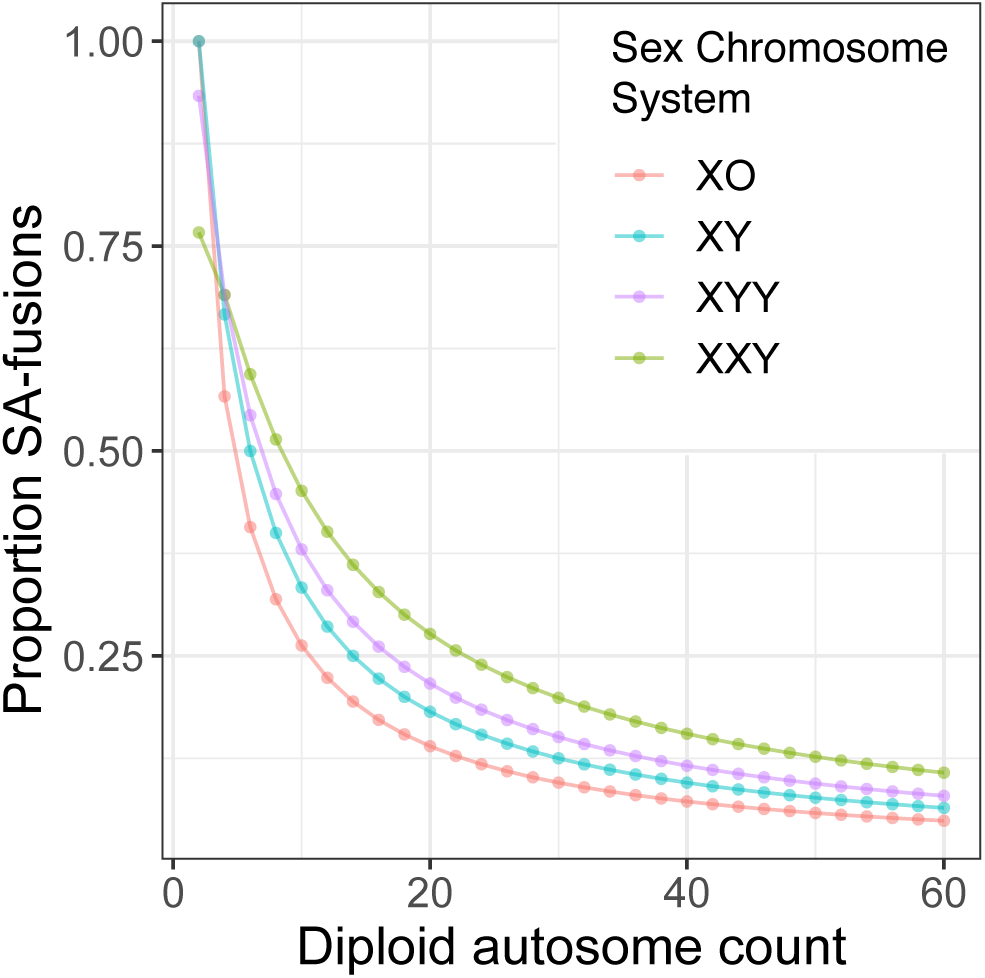
Probability of a random fusion joining a sex chromosome and autosome. On the vertical axis we plot the proportion of all fusions that are SA-fusions while on the horizontal axis we plot the diploid autosome count. Each sex chromosome system is indicated by a unique color.

In the previous example, we calculated the expected proportion of the different types of fusions based on the ancestral, and most common, karyotype inferred in a clade. However across the entire clade, a variety of karyotypes exist. We envision the primary use of equation 11 will be to calculate the expected proportion of fusions that are SA-fusions across large clades. We can do this by employing a biologically realistic Markov model of possible fusions and fissions (Blackmon et al. 2019), and leveraging stochastic mappings generated under such a model to extract the proportion of time that lineages in a clade spent with each possible chromosome number and sex chromosome system (Huelsenbeck et al. 2003; Revell 2012). These proportions can then be used in conjunction with equation 11 to generate a weighted sum that describes the expected proportion of all observed fusions that are SA-fusions (figure 2). The resulting expected value can then be compared to the observed proportion of SA-fusions inferred from the stochastic mappings. An additional advantage of this approach is that it naturally extends to marginalize over a collection of phylogenetic trees sampled from a posterior distribution. This approach would pro

**Figure 2:**
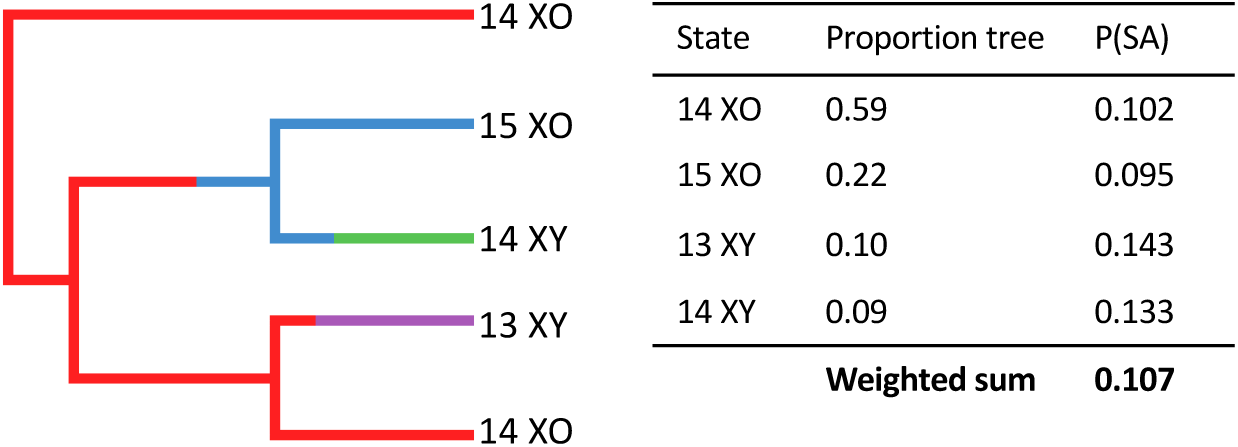
Estimating *P*(*SA*) across a clade. On the left a stochastic map showing chromosome number and sex chromosome system. In the table on the right we have calculated the proportion of time that each state is present in the clade and then calculated *P*(*SA*) for each of these states. These *P*(*SA*) values along with the proportions are used to generate the expected *P*(*SA*) for the clade as a whole.

We have developed a flexible equation used to calculate the probability of SA-fusions under common sex chromosome systems (male or female heterogametic). This model will allow for quantitative analyses of fusions across large clades and provide a way to test the long-standing hypothesis that SA-fusions are selectively favored for their ability to resolve sexual antagonism. In some clades where chromosome number is high (e.g. Lepidoptera and Isoptera) our model shows that SA-fusions should be rare (Blackmon et al. 2017). In these cases, several SA-fusions within a clade may well suggest that these fusions are selectively favored. However, this model also shows that for clades with very few chromosomes (e.g. Diptera and Hemiptera), we should expect many SA-fusions even if they are not selectively favored (Blackmon et al. 2017). Therefore, SA-fusions should only be considered as evidence for sexual antagonism when they occur at a higher rate than expected for the chromosome numbers and sex chromosome systems that have been present during the evolution of a clade.

## Literature Cited

Ashman, T.-L., D. Bachtrog, H. Blackmon, E. E. Goldberg, M. W. Hahn, M. Kirkpatrick, J. Kitano, J. E. Mank, I. Mayrose, R. Ming, et al., 2014. Tree of sex: A database of sexual systems. Scientific Data 1:140015.

Bachtrog, D., J. E. Mank, C. L. Peichel, M. Kirkpatrick, S. P. Otto, T.-L. Ashman, M. W. Hahn, J. Kitano, I. Mayrose, R. Ming, et al., 2014. Sex determination: why so many ways of doing it? PLoS Biol 12:e1001899.

Blackmon, H., J. Justison, I. Mayrose, and E. E. Goldberg, 2019. Meiotic drive shapes rates of karyotype evolution in mammals. Evolution 73:511–523.

Blackmon, H., L. Ross, and D. Bachtrog, 2017. Sex determination, sex chromosomes, and karyotype evolution in insects. Journal of Heredity 108:78–93.

Charlesworth, D. and B. Charlesworth, 1980. Sex differences in fitness and selection for centric fusions between sex-chromosomes and autosomes. Genetics Research 35:205–214.

Cheng, C. and M. Kirkpatrick, 2016. Sex-specific selection and sex-biased gene expression in humans and flies. PLoS Genetics 12.

Hsu, T. C. and K. Benirschke, 2013. An atlas of mammalian chromosomes, vol. 10. Springer Science & Business Media.

Huelsenbeck, J. P., R. Nielsen, and J. P. Bollback, 2003. Stochastic mapping of morphological characters. Systematic Biology 52:131–158.

Innocenti, P. and E. H. Morrow, 2010. The sexually antagonistic genes of drosophila melanogaster. PLoS biology 8.

Kasimatis, K. R., P. L. Ralph, and P. C. Phillips, 2019. Limits to genomic divergence under sexually antagonistic selection. G3: Genes, Genomes, Genetics 9:3813–3824.

Maddison, W. P. and G. Leduc-Robert, 2013. Multiple origins of sex chromosome fusions correlated with chiasma localization in habronattus jumping spiders (araneae: Salticidae). Evolution 67:2258–2272.

McAllister, B. F., 2003. Sequence differentiation associated with an inversion on the neo-x chromosome of drosophila americana. Genetics Society of America 165:1317–1328.

Pennell, M. W., M. Kirkpatrick, S. P. Otto, J. C. Vamosi, C. L. Peichel, N. Valenzuela, and J. Ki- tano, 2015. Y fuse? sex chromosome fusions in fishes and reptiles. PLoS genetics 11.

Ponnikas, S., H. Sigeman, J. K. Abbott, and B. Hansson, 2018. Why do sex chromosomes stop recombining? Trends in Genetics 34:492–503.

Revell, L. J., 2012. phytools: an r package for phylogenetic comparative biology (and other things). Methods in ecology and evolution 3:217–223.

Stebbins, G. L. et al., 1971. Chromosomal evolution in higher plants. Chromosomal evolution in higher plants..

White, M. J. D., 1977. Animal cytology and evolution. CUP Archive.

Zhou, Q. and D. Bachtrog, 2012. Sex-specific adaptation drives early sex chromosome evolution in drosophila. Science 337:341–345.

