## supplemental file 1 for "The Probability of Fusions Joining Sex Chromosomes and Autosomes": supplementalfile1.R

```
# Nathan W. Anderson and Heath Blackmon
# 27 March 2020
#

# Each of the three functions below contains three arguments
# Da: diploid count of autosomes
# scs: scs system possible values are
#      XO, XXO, XY, XYY, XXY, ZO, ZZO, ZW, ZWW, ZZW
# mud: proportion of fusions that originate in females
#      should be 0.5 unless other data is avaialable

# function for P(SA)
Pfsa <- function(Da, scs, mud = 0.5){
  sexchroms <- strsplit(scs, split = "")[[1]]
  if(scs %in% c("XO", "XXO", "XY", "XYY", "XXY", "XYYY", "XXXY")){
    Xs <- sum(sexchroms == "X")
    Y <- sum(sexchroms == "Y")
    Ds <- Da + Xs + Y
    Dd <- Da + 2 * Xs
    res <- 1 - mud * ((Da * (Da - 2) + 2 * Xs * (2 * Xs - 2)) / (Dd * (Dd - 2))) -
      (1 - mud) * ((Da * (Da - 2) + max(c(Xs, Y)) * (max(c(Xs, Y)) - 1)) / (Ds * (Ds - 2)))
  }
  if(scs %in% c("ZO", "ZZO", "ZW", "ZWW", "ZZW", "ZWWW", "ZZZW")){
    Zd <- sum(sexchroms == "Z")
    W <- sum(sexchroms == "W")
    Dd <- Da + Zd + W
    Ds <- Da + 2 * Zd
    mus <- 1-mud
    res <- 1 - mus * ((Da * (Da - 2) + 2 * Zd * (2 * Zd - 2)) / (2 * Ds * (Ds - 2))) -
      (1 - mus) * ((Da * (Da - 2) + max(c(Zd, W)) * (max(c(Zd, W)) - 1)) / (2 * Dd * (Dd - 2)))
  }
  return(res)
}
# function for P(AA)
Pfaa <- function(Da, scs, mud = 0.5){
  sexchroms <- strsplit(scs, split = "")[[1]]
  if(scs %in% c("XO", "XXO", "XY", "XYY", "XXY", "XYYY", "XXXY")){
    sexchroms <- strsplit(scs, split="")[[1]]
    Xs <- sum(sexchroms == "X")
    Y <- sum(sexchroms == "Y")
    Ds <- Da + Xs + Y
    Dd <- Da + 2 * Xs
    res <- mud * ((Da * (Da - 2)) / (Dd * (Dd - 2))) +
      (1 - mud) * ((Da * (Da - 2)) / (Ds * (Ds - 2)))
  }
  if(scs %in% c("ZO", "ZZO", "ZW", "ZWW", "ZZW", "ZWWW", "ZZZW")){
    Zd <- sum(sexchroms == "Z")
    W <- sum(sexchroms == "W")
    Dd <- Da + Zd + W
    Ds <- Da + 2 * Zd
    mus <- 1-mud
    res <- mus * ((Da * (Da - 2)) / (Ds * (Ds - 2))) +
      (1 - mus) * ((Da * (Da - 2)) / (Dd * (Dd - 2)))
  }
  return(res)
}
# function for P(SS)
Pfss <- function(Da, scs, mud = 0.5){
  sexchroms <- strsplit(scs, split = "")[[1]]
  if(scs %in% c("XO", "XXO", "XY", "XYY", "XXY", "XYYY", "XXXY")){
    Xs <- sum(sexchroms == "X")
    Y <- sum(sexchroms == "Y")
    Ds <- Da + Xs + Y
    Dd <- Da + 2 * Xs
    res <- mud * ((2 * Xs * (2 * Xs - 2)) / (Dd * (Dd - 2))) +
      (1 - mud) * ((max(Xs, Y) * (max(Xs, Y) - 1))/(Ds * (Ds - 2)))
  }
  if(scs %in% c("ZO", "ZZO", "ZW", "ZWW", "ZZW", "ZWWW", "ZZZW")){
    Zd <- sum(sexchroms == "Z")
    W <- sum(sexchroms == "W")
    Dd <- Da + Zd + W
    Ds <- Da + 2 * Zd
    mus <- 1 - mud
    res <- mus * ((2 * Zd * (2 * Zd - 2))/(Ds * (Ds - 2))) +
      (1 - mus) * ((max(Zd, W) * (max(Zd, W) - 1))/(Dd * (Dd - 2)))
  }
  return(res)
}

# calculating pval from line 194
sum = 0
for(i in 8:10){
  for(j in 0:(10-i)){
    sum <- sum + (factorial(10)/(factorial(i)*factorial(j)*factorial(10-i-j))) *
      Pfsa(26, 'XXO', .5)^i * Pfaa(26, 'XXO', .5)^j * Pfss(26, 'XXO', .5)^(10-i-j)
  }
}
p.val <- sum
p.val
```
